# Strategic use of male alternative reproductive tactics in cooperatively breeding banded mongoose groups

**DOI:** 10.1101/2024.03.11.584460

**Authors:** Graham Birch, Hazel J. Nichols, Francis Mwanguhya, Jonathan D. Blount, Michael A. Cant

**Author notes:** Authors for correspondence G. Birch ( /).

## Abstract

Alternative reproductive tactics (ARTs) allow smaller or less competitive individuals to reproduce by avoiding direct fights through sneaky strategies. Within cooperatively breeding groups ARTs are rarely reported, potentially due to difficulties observing male reproductive behaviour in the wild, or reproductive suppression by fellow group members. In societies where all mating opportunities cannot be monopolised by one male, young males could use sneaky tactics as a ‘stepping-stone’ to gain limited reproductive success while they grow in resource holding potential (RHP). Using 20 years of pedigree, weight, group demography, and behavioural data, we investigated the use of sneaky ‘pesterer’ ARTs in wild banded mongooses. Instead of a ‘stepping stone’, pestering tactics were typically a rare back-up used by males displaced from mate guarding tactics by higher RHP rivals. Additionally, pesterers have lower siring success compared to mate guards, despite similar weight loss costs, which may explain why most males settle for reproductive inactivity rather than use pestering tactics. However, pestering allows younger males to gain access to older, higher fecundity females in the group and may even facilitate inbreeding avoidance. Overall, ARTs provide a viable option for reproductive fitness for some males that lose out on reproductive positions in highly male-skewed groups.

## Introduction

In animal societies, male reproductive success is typically skewed towards those males with the highest resource holding potential (RHP)[1,2], defined as a measure of “absolute fighting ability” [3]. These high RHP males are able to monopolise the best territories [4,5], more successfully guard females [6,7] or defend resources that attract females [8,9]. A high RHP is associated with prime age classes [4,10], a large body size [11], or effective weaponry [12], all of which provide an advantage competing for mates. Where high RHP individuals can outcompete other males through fights, there may be selection for low RHP individuals to adopt sneaky alternative reproductive tactics (ARTs)[13]. ARTs are discrete reproductive phenotypes present in the same population [14]. Compared to fighters who directly compete to monopolise access to mates, ARTs often employ parasitic or sneaky tactics. Sneaky tactics can give lower RHP males opportunities for reproductive success through means that avoid the costs of direct confrontation with superior rivals [15–21].

ARTs are taxonomically widespread and take many forms including differences in behaviour or morphology [14,22]. Sneaky morphotypes can be fixed from birth by genes [13,23–25], such as small and large morphs of the pygmy sword tail fish *Xiphophorus nigrensis* [26], from parental effects such as in spider mites *Tetranychus urticae* [18], or by the developmental environment [27–29] such as the effect of temperature on growth and ART expression in the male squid *Uroteuthis edulis* [19]. Fixed ARTs are typically under negative frequency dependent dynamics[30–32], meaning that they are more successful when rare in a population. Good evidence in support of negative frequency dependent dynamics is found in male blue gill sunfish (*Lepomis macrochirus*) where the fitness payoff of sneaking morphs decreases as they became more common [33].

In contrast to fixed morphologies many ARTs, including sneaky tactics, exhibit flexibility [30,34,35]. This flexbility should allow individuals to strategically choose the best suited tactic given their relative success in the current environment. For example, the success of two distinct sneaky ARTs in Ornate tree lizards (*Urosaurus ornatus)* is determined by food availability triggering adaptive tactic switching in males [36]. Flexible use of sneaky tactics is often described as a ‘best of a bad job’, used by lower RHP males in the population who are at a competitive disadvantage who would not have success fighting directly with rivals [30,37,38]. By opening up opportunities when at a competitive disadvantage, the strategic adoption of sneaky ARTs could increase life-time reproductive fitness [39,40].

One strategic use of sneaky tactics is as a conditional ‘stepping stone’. In this pattern of alternative reproductive tactic (ART) adoption, young males with lower RHP initially employ sneaky tactics and transition to fighting tactics only when they have grown sufficiently to compete successfully with rivals [30,31,41–43]. In the cichlid *Lamprologus callipterus,* small sneakers as they grow eventually become territorial nest builders to attract females themselves [44,45]. Similarly, young African striped mice *(Rhabdomys pumilio)* may use roaming tactics until they grow large enough to defend a harem of females [41,42]. These ’stepping stone’ dynamics may be driven by reproductive costs, as lower RHP males are more prone to injuries in fights and may lack the condition required to sustain the energetic demands of reproductive activity compared to their higher RHP rivals [46,47]. Sneaky tactics may avoid these costs, such as in the two-spotted spider mite (*Tetranychus urticae Koch*) where young males increase their survival when using sneaky as opposed to fighting reproductive tactics [48]. The use of sneaky tactics as a stepping stone allows males to gain early reproductive success by avoiding the costs of fighting superior rivals, while positioning themselves for a future transition to more profitable fighting tactics.

Another strategic use of ARTs is a back-up option when low RHP fighters are displaced from reproductive positions. Sneaky ARTs may allow males to recover access to reproductive opportunities when they are displaced entirety from reproductive positions. For example, in Seba’s short tailed bats, *Carollia perspicillata,* males who defended their own harems in the past may settle for peripheral roles as they senesce, suggesting these old males utilise sneaky ARTs as a back-up option for reproductive success after displacement by rivals [31]. Males may also be displaced from higher to lower quality mating opportunities, such as from high to low fecundity females [49], but these males regain access by using sneaky ARTs. Overall, sneaky ARTs may provide a back-up option to increase their reproductive fitness when attempts to fight fail due to competition.

Here we describe and evaluate the strategic use and adoption behind a unique sneaky ART in cooperative breeding banded mongoose (*Mungus mungo)* groups. Male competition is highly costly in cooperative breeding groups with fatalities in fights common [1,50,51]. In many cooperative breeding groups a dominant pair typically supress the reproduction of other mature group members [52–54], with maturing males often forced to disperse to reproduce [2,55], or wait to inherit breeding positions [56]. Instead, in banded mongooses secondary reproducing males that use sneaky ARTs are tolerated in intermediate reproductive skew groups where multiple males and females breed. Banded mongooses groups are highly male skewed, typically with multiple mature males for every breeding female [57]. Dominant ‘guarding’ males can only monopolise reproduction from one female at a time over the duration of short synchronrous group oestrus events. Guards follow females continuously while aggressively preventing copulations from rivals [58]. Similar guarding and territorial activity has been associated with costly energetic demands [59,60], and males may suffer opportunity costs from reduced forraging time. Guarding follows assortative mating patterns, with the oldest males guarding the oldest most fecund females, resulting in the top three males siring the majority of the group’s offspring [61]. ‘Pesterers’ attempt to sneak copulations when guards become distracted or during attempts by females to escape their guard [58]. Many other mature males are reproductively inactive, displaying neither guarding nor pestering behaviour [57,61]. Our primary aim was to test the hypothesis that males use sneaky tactics as a ‘stepping stone’ strategy, to gain matings while waiting to grow in size, RHP and experience. We also compared the relative costs and fitness benefits associated with mate-guarding versus sneaky tactics.

Our secondary aim was to investigate the hypothesis that the use of sneaky ARTs offers a strategic advantage to males in terms of mate choice. For male mate choice to evolve, males should be forced into time-limited simultaneous assessments over females that vary in aspects of quality such as fecundity [62,63]. To make the best use of their time, high RHP fighters should pursue the most in-demand fecund females, displacing low RHP rivals to less fecund females. The patterns of mate choice that result where the quality of males and females is matched is known as assortative mating [49]. Low RHP males, by sneaking, could gain access to higher quality females than they could if they tried to monopolise them directly. Furthermore, a second assessment for mate choice is genetic compatibility [64]. Like most cooperative breeders, members of banded mongoose groups are typically closely related, and support for mate choice by mate guards and reproducing females to avoid inbreeding depression has been found in the past [65,66]. If sneaky ARTs open up male mate choice options they may give an advantage accessing less related females and avoiding inbreeding. ARTs may be favoured by females who gain access to males they would otherwise be prevented from mating with, as suggested by previous observations of females attempting to escape their mate guards[58].

Our analysis proceeds in three steps. First, we assess the relative costs and fitness benefits of ARTs in male banded mongooses by comparing weight loss and siring success between mate guarding and sneaky pestering tactics. As a ‘best of a bad job’ tactic we expect sneakers to have less access to copulations than guards, hence we predict that sneakers will have reduced siring success compared to guards. According to negative frequency dependent dynamics, we also predict pestering ARTs will have a higher payoff when rare.

Second we assess why males might favour switching to pestering tactics by comparing tactic transitions of inactive and reproductively active males. Specifically, we will test whether males transition from reproductive activity to pestering as a stepping stone on the way to a guarding tactic.

Third, we assess whether pestering tactics provide males with increased opportunities for mate choice. We predict that mate guards will be more restricted in their access to mates, and guard a more limited pool of females, compared with those that pestering males pursue. We also test whether sneaky males have access to less closely related females, suggesting that this tactic may yield benefits to both males and females in terms of inbreeding avoidance.

## Methods

### Study population

The banded mongoose study population is located on Mweya Peninsula, Queen Elizabeth National Park, Uganda (0°12′S, 29°54′E). The history of each individual and group is known through life-history data collection ongoing since 1995 [67]. For identification, individuals are given a unique fur shave patterns on their back. Data used in this study is from the period between April 2003 and February 2021.

### Behavioural data collection

#### Male behaviour monitoring during group oestrus events

Groups were visited at least every 3 days to collect life-history data. Where signs of oestrus were found, groups were visited every day until reproductive behaviour ceased. Each day during oestrus every breeding female received independent observation (focals) lasting 20 minutes. During focals, any males that guarded or pestered the focal female were noted. Guards were identified as a single male that followed the focal female within 5 metres throughout the 20 minutes, and pesterers as any male that tried to interrupt the focal-guard pair or make attempts to copulate with the focal female if the guard was distracted. Focals took place during peak foraging periods in the morning, and were paused where view of the focal female was obscured or during group alarms. Groups continued to be visited every day to collect focals until reproductive behaviour ceased. To avoid sampling immature males, data on males younger than subadults (less than 180 days of age) was discounted in all analyses

Most oestrus events (events spanning the start of oestrus until reproductive behaviours cease) had multiple days of data collection (n= 379, mean:2.997, IQR:3). On each day males were noted as a guard or pesterer according to their behaviour towards females, and as a inactive if they showed no interest. For the purposes of examining behaviour transitions between oestrus events, male reproductive behaviour from all data collection days was summarised into one state for each oestrus event. If males guarded on 50% or more data collection days, such individuals were defined as a guard, while males were assigned as subordinates if they were inactive for the whole oestrus event. Males that did not guard on 50% or more days, but instead pestered on more than 50% of data collection days, were defined as a pesterer.

### Age, weight and demographic data

All males in this analysis were followed from birth to death and are therefore of a known age. Many males share age ranks as demography of groups is stratified around surviving siblings of shared litters. In our analysis shared age ranks were assigned to the minimum rank and subsequent younger males ranks left a gap, for example if there were 3 age rank 2 males the next oldest was assigned age rank 5.

Body weights have been collected for the population since 2000. Weight collection frequency varies between individuals and groups depending on their degree of habituation. To account for this variation, weights were averaged over a shared time period. For the state transition and paternity analysis, weights were averaged for the 60 days before and after a given oestrus event (oestrus weight). To control for variation in male weights between groups we centred weight around the group mean (individual oestrus weight – mean male oestrus weight). 6.6% (239/3638) of oestrus weights could not be calculated with data available. Imputation has been carried out in previous studies on long term populations where missing data can occur [68,69], including this same population of banded mongooses [70]. Using the full history of weight collection for each individual, and the correlation between age and weight in banded mongooses, these missing weights were inputted (see supplementary for more detail).

Predicting the likely weight of males through imputations methods is not applicable to measurement of actual weight loss, therefore for weight loss models prior and post oestrus weights were extracted separately. Prior oestrus weights were averaged over the 60 days prior to each oestrus event and the post oestrus weight was the nearest single weight recorded after an oestrus event ceased. A preliminary model was run to detect an effect of time to post-oestrus weight collection on weight loss, of which no significant effect was found when restricting sampling to one week post-oestrus. Males with missing weights were simply not used.

### Sired litters and group-centered relatedness

A genetic pedigree has been collected since 2003 [see references for how the pedigree is obtained ;11,40]). The methods for constructing the genetic pedigree and calculating pair-wise relatedness for the banded mongoose population have been described previously [71]. A banded mongoose’s gestation period is around 9 weeks [pooled data from 58,65]. 57 litters could be connected to 35 mothers that received attention from at least one pestering and guarding male in oestrus events approxiamtely 9 weeks prior to the litters birth date (ended 59 ± 15 days before). This included 60 cases of succesful sires (3 had multiple paternity) from interactions with 104 unique males.

Group-centered relatedness was calculated by taking the average pedigree relatedness of a given male to every adult female present in the same group during an oestrus event, and then for each male the above average was deducted from pedigree relatedness to each adult female present in the group.

### Statistical analysis

All models were fit using Bayesian inference (JAGS MCMC) in R[72,73]. To improve model convergence, numeric covariates with a range below 0 and above 1 were standardised. Multicolinearity was checked using the crosscorelation plots using the ggmcmc package[74]. Chain convergence was checked using rhat values from the JAGS model output, with all models showing convergence of chains for each fitted parameter (R<1.1). Convergence was also checked using traceplots by eye (ggmcmc).

### State transition modelling

The transition probability of males from one reproductive status to the next was modelled using JAGS MCMC [31,72]. This model was based on 3638 state transitions of which 639 involved transitions to or from a pestering tactic. The model used state matrix **z** with element **z***_i,t_* for the state of male *i* at oestrus event *t. i* comprises the 320 males that have been followed from birth to death in the study population. *t* comprises each oestrus event in order a given male has participated in throughout their life. *A* state-transition matrix **Ω** was set with four dimensions, previous reproductive state *n*, new state *m*, male *i,* and oestrus event *t*. The state process **ω***_n,m,I,t_* represents the probability that male at reproductive state *n* at oestrus event *t*, will be in state *m* at the next oestrus event *t+1*. The probability of leaving any reproductive status and dying before the next oestrus event was defined as 1 minus the reproductive status related survival probability. Once a male had died its probability of remaining dead was defined as 1. Since we have complete records of group composition during oestrus events, therefore each male *i* had reproductive state data for each oestrus event *t* they lived through, an observation matrix was not required (always observed). Using the state-transition matrix **Ω** reproductive state transitions (Iterations=20000, Thinning interval =100, burn in=2500, Chains=3) were regressed against age and weight relative to other males in the group (age rank and group centred weight) as a metric of RHP (metrics of RHP). Running separate models for moderately correlated variables may lead to bias exaggerating their significance, justifying age rank and group centered weights (r=0.44) inclusion in the same model [75]. Interactions between group centered weight and age rank were however not fitted as issues with multicollinearity would be exaggerated [75]. Group sex ratio was fitted in the same model (number of males over number of females of at-least one year of age) to test for an effect of competition. Interactions between group sex ratio and both age rank and group centered weight were fitted but were removed when interaction terms proved not credible. To control for common group membership during oestrus events and repeated sampling on the same males, random effects for oestrus event ID (n=375), group ID (n=20), and male ID (n=320) were fitted.

### Weight loss models

554 weight changes from subordinates, 170 weight changes from pesterers, and 265 weight changes from guards were included. To control for common group membership during oestrus events and repeated sampling on the same males, random effects for oestrus event ID (n=118), historical group ID (n=12), and male ID (n=259) were fitted. Percentage weight loss for each individual over an oestrus event was normally distributed, and as such was regressed with behavioural state (subordinate versus guard) using a Gaussian distribution in JAGS mcmc (Iterations=50000, Thinning interval =100, burn in=5000, Chains=3).

### Bernoulli models for mate choice and fitness

There were 57 cases where a litter could be attributed with high confidence to an oestrus event where the female was observed to be guarded and pestered by different males, which included the sire verified through pedigree data taken from 2003-2019, see [61,76] for methodology. This included 104 unique males that interacted with 35 unique females in 41 oestrus events, included as random effects. For each dyadic interaction, a binomial success or fail was determined depending on whether the male succeeded in siring offspring. The probability that a given male would successfully sire was modelled using Bernoulli GLMM fitted in JAGS MCMC (iterations=100000, thinning interval =100, burn in=10000, chains=3). The model was fitted with the reproductive state of each male to test whether guards were more successful than pesterers (guards as 1, pesterers as 0), and proportion of competitors adopting the same tactic who interacted with the same female. Guards and pesterers were defined according to their average behaviour towards a given female overall in the oestrus event as in other analyses (if guarded >50% of days then guard, ties=guard, pestered>50% of days then pesterer). To control for the fact that each new male interacting with a female decreased the chance each male had of siring her offspring, the number of male competitors interacting with each female over the course of the oestrus event was fitted as a fixed effect. To assess if the effect of reproductive state was influenced by its commonality, interaction terms were initially included, but dropped from the final model when they were not proved credible.

For each day where a male over 180 days of age has interacted (as a guard or pesterer), a binomial variable was added for each female present in the group to distinguish the specific females the male interacted with. This included 1558 guarding and 674 pestering interactions involving 228 unique adult males and 230 unique adult females in 297 oestrus events. The smaller number of oestrus events and males compared to the state transition analysis is due to only including interactions where the relatedness of the male and female were known, as well as the age of the female involved. Only rarely did a given male interact with multiple females on the same day (308/2232 ∼14% of cases), yet as different groups of females enter and leave oestrus across multiple days males commonly (927/1298 ∼71.4% of cases) interacted with multiple potential mates over the course of the oestrus event. The probability that a given male would interact with a female was modelled using a Bernoulli GLMM fitted in JAGS MCMC (iterations=20000, thinning interval =100, burn in=2000, chains=3). This was regressed with group centered relatedness, and an interaction between male and female age rank to assess for patterns of assortative mating in banded mongooses. Further, interactions with reproductive state were fitted, a three-way interaction with male and female age rank and two-way with relatedness, respectively, to assess for whether guards and pesterers have different preferences for females. The interaction between relatedness and state was dropped when it did not prove credible. To account for the probability of interacting being inversely associated with the number of females, the number of adult females present in the group was added as a fixed covariate.

### Ethics

Prior approval of all work was received from Uganda Wildlife Authority (UWA) and Uganda National Council for Science and Technology (UNCST). The Ethical Review Committee of the University of XXXXX approved all research activites.

## Results

Pesterers lost a similar amount of weight (Supplementary figure S1; mean=-2.3% bodyweight, hci=-3.37%, lci=-1.28%) to guards (mean=-2.31%, hci=-1.47%, lci=-3.21%) over the course of an oestrus event, over subordinates that had a statistically neutral weight change (mean=0.32%, hci=1.11, lci=-0.48).

Although on each day a given male is usually recorded interacting with a single female, most males interacted with multiple females over the course of the oestrus event, with pesterers on average interacting with more females (mean females interacted with per day of oestrus: guards = 1.11 ±0.01s.e.m, pesterers=1.18±0.02 2s.e.m, and mean females interacted with over each oestrus event: guards=2.34 ±0.052s.e.m, pesterers= 2.71±0.05s.e.m).

Guards had a significantly higher (mean=0.4, hci=0.52, lci=0.28) chance than pestering competitors (mean=0.12, hci=0.19, lci=0.6) to sire a female’s offspring (supplementary table S1a). There was no significant interaction between the number of competitors and tactic (guard versus pesterer) on success at siring offspring, with guard maintaining a siring advantage against single pesterers (supplementary figure S2 - guard: mean=0.65, hci=0.80, lci=0.46 vs pesterer: mean=0.27, hci=0.42, lci=0.14) or multiple pesterers, e.g. 2 (guard: mean=0.67, hci=0.83, lci=0.47 versus pesterers: mean=0.13, hci=0.54, lci=0.23).

There was a significant negative effect of the commonness of a reproductive tactic among males interacting with the same female on the chance to sire offspring. However, there was no evidence of an interaction with tactic, suggesting the success of pestering and guarding males are under similar negative frequency dependent dynamics. For example, when controlling for competition and given a scenario where 3 males interact with the same female, independent of the tactic adopted, the 2 males adopting the same tactic have a relative disadvantage (mean=0.26, hci=0.4, lci=0.15) to the lone male adopting the rarer tactic (mean=0.48, hci=0.66, lci=0.3).

Transitions to pestering tactics were rare overall (Figure 1ai,bi,ci). On average, inactive subordinates transitioned into guarding tactics (mean=0.2, hci=0.28, lci=0.16) more often than into pestering tactics (mean=0.08, hci=0.09, lci=0.07). Pestering was the rarest transition from a guarding tactic (mean=0.13, hci=0.15, lci=0.1), with keeping a guarding tactic (mean=0.41, hci=0.5, lci=0.34) statistically similar to dropping into a subordinate tactic (mean=0.46, hci=0.52, lci=0.39). Pestering tactics themselves were unstable having a lower probability of being retained (mean=0.17, hci=0.21, lci=0.12) than gaining a guarding tactic (mean=0.35, hci=0.45, lci=0.25), with dropping into a subordinate tactic being the most likely transition (mean=0.49, hci=0.57, lci=0.4). Transition probabilities from a pestering tactic resemble transition probabilities from a guarding tactic (Figure 1b,1c), suggesting pesterers and guards are functionally similar in their future use of tactics.

**Figure 1:**
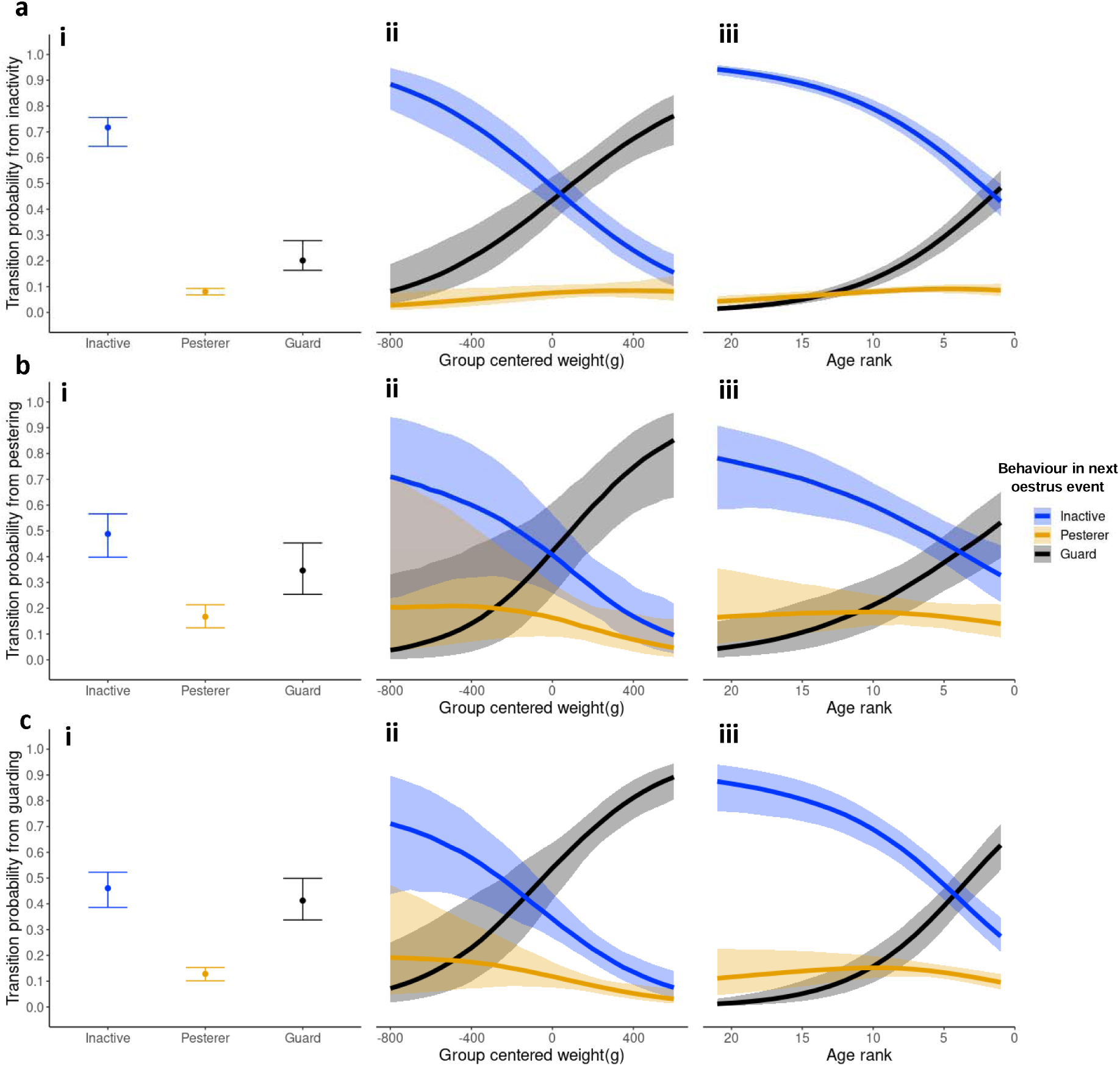
The effect of age rank and group-centered weight on between-oestrus event state transition probabilities. Lines represent the mean posterior probability and ribbons are 95% credible intervals. Each panel represent transition probabilities from the 3 distinct states. Black lines and ribbons correspond to transitions to guarding, orange transitions to pestering, and blue transitions to subordinate tactics. Panel a) shows mean (points) and credible intervals (error bars) for all 9 transitions for reference based of a null model with no covariates.

Age rank and group centred weight were correlated (r=-0.44), and the effect of increasing weight on transition probabilities largely mirrored the effect of decreasing age rank (Figure 1ii, iii). In all reproductive states there was a negative effect of age rank (older) on transitions to subordinate, and positive (younger) to a guarding tactic (supplementary table S1a). Therefore, as males aged they became more likely to adopt a future guarding tactic and less likely to adopt a subordinate tactic (Figure 1iii). Similaly, there was a significant negative effect of weight on transitions to subordinate, and a positive effect of weight on transitions into guarding tactics (Figure 1ii). As males moved into older age rank and grew in weight, their probability to transition into a pestering tactic from all reproductive states significantly decreased (Supplementary table S1a), with the exception of weight having no significant effect on subordinate to pestering transition probabilities. Posterior probabilities to transition to a pestering tactic (figure 1) only started to decline from a peak at intermediate age ranks (from subordinate: age rank 5, mean=0.09, hci=0.11, lci=0.08, from pesterer: age rank 11, mean=0.19, hci=0.26, lci=0.23, from guard: age rank 10, mean=0.15, hci=0.2, lci=0.11) and weights (from pesterer: -250g, mean=0.19, hci=0.3, lci=0.12, from guard: -200g, mean=0.16, hci=0.23, lci=0.1), continuing to decline towards age rank 1 (from subordinate: mean=0.09, hci=0.11, lci=0.06, from pesterer: mean=0.14, hci=0.21, lci=0.09, from guard: mean=0.1, hci=0.13, lci=0.07) and the heaviest weights e.g +400g (from pesterer: mean=0.09, hci=0.18, lci=0.04, from guard: mean=0.06, hci=0.1, lci=0.03).

Remaining or becoming a subordinate dominated transitions for young males, but if young subordinates (below age rank 19) did become active they transitioned into pestering tactics more often than to guarding tactics (to pesterer: mean=0.05, hci=0.07, lci=0.03, to guard: age rank 20, mean=0.02, hci=0.03, lci=0.01). In comparison, slightly older guards, younger than age rank 17, were more like to transition into a pestering tactic than to keep their guarding tactic (age rank 18 to pesterer: mean=0.13, hci=0.22, lci=0.06, to guard: age rank 20, mean=0.03, hci=0.06, lci=0.01). As individuals move into older age ranks, transitions to a guarding and pestering tactic became statistically similar until guarding transitions overtook pestering transition probabilities at age rank 9 in inactive subordinates (to pesterer: mean=0.08, hci=0.09, lci=0.07, to guard: mean=0.13, hci=0.16, lci=0.1), and at age rank 7 in guards (to pesterer: mean=0.15, hci=0.18, lci=0.12, to guard: mean=0.28, hci=0.36, lci=0.2).

Sex ratios were on average male biased (mean=2.2 males per female IQR=1.4). There was no significant effect of biased sex ratios on any tactic transition probabilities (see supplementary for more detail, table S1a, Figure S3).

There was a significant interaction between state (guard or pesterer), and male and female age rank on the probability a male interacted with a given female (supplementary table S1d). Males at the oldest age ranks (1-3) typically interacted with the oldest females more often regardless of their reproductive tactic. For example, given 7 females in the group, guards and pesterers interact with age rank 1 females around a third of the time (Figure 2: state=guard, mean= 0.33, hci=0.36, lci=0.3 vs state=pesterer, mean=0.34, hci=0.39, lci=0.29), compared to age rank 7 females around 1 in 10 times (state=guard, mean= 0.08, hci=0.09, lci=0.06 vs state=pesterer, mean=0.1, hci=0.12, lci=0.08). Guarding and pestering interactions began to diverge at younger age ranks. Guards continued to assortatively mate, with the youngest males interacting with the youngest females more often. For example, if there are 7 females in the group, males of age rank 10-12 were far more likely to interact with age rank 7 females (mean=0.28, hci-0.32, lci=0.24) than the top females (i.e. age rank 1:, mean=0.1, hci=0.12, lci=0.08). Although younger pesterers did interact with the top females less than older pesterers, this switch in interaction probability towards younger females noticeably lagged behind guards (Figure 2), with the youngest pesterers such as age rank 10-12 continuing to interact with the top females as often as the youngest females (age rank 1 females:, mean=0.17, hci=0.21, lci=0.14 versus age rank 7 females: mean=0.17, hci=0.21, lci=0.14). This suggests pesterers interacted with older females more than would be expected if they adopted a guarding tactic.

**Figure 2:**
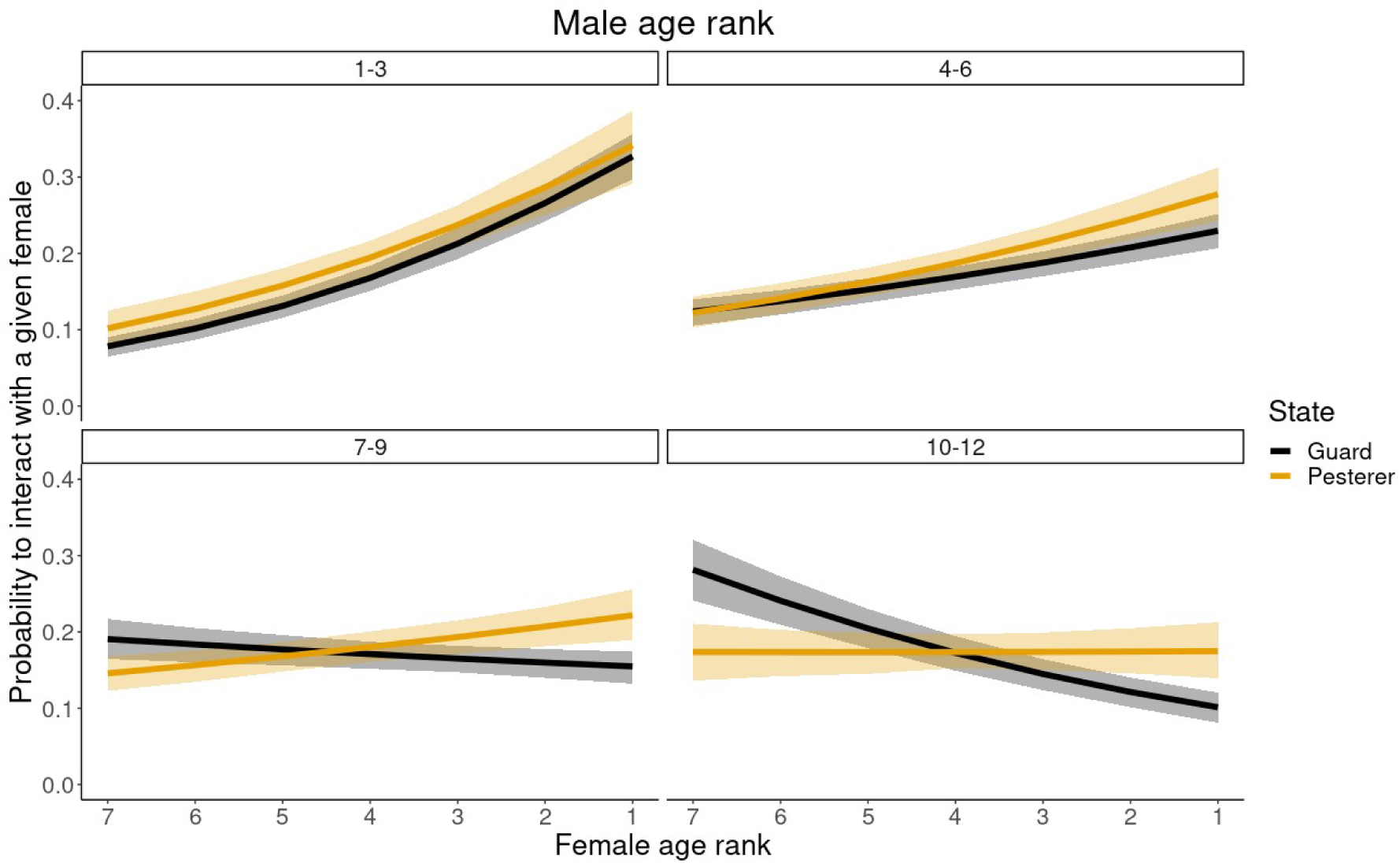
Probability that males interact with females of different age rank, for four different age categories of male, from oldest ranked males (top left) to youngest (bottom right). Black lines are guards, orange lines are pesterers. Lines represent the mean posterior probability and ribbons areb 95% credible intervals. Posteriors probabilities have been averaged for age rank bands of 3 for plotting purposes. To control for amount of choice, these probabilities are modelled given there are 7 females present in the group.

Males had a preference for less closely related females compared to females with a high relatedness to themselves (supplementary table S1 e.g. group centered relatedness = -0.25, mean given 7 females present=0.21, hci=0.24, lci=0.19; group centered relatedness =0.35: mean given 7 females present=0.15, hci=0.17, lci=0.13). There was no interaction between reproductive state and relatedness, suggesting avoidance of inbreeding does not differ amongst guards and pesterers. There was also no interaction between a male’s age rank a female’s relatedness to interacting males, suggesting younger males were not restricted in their choice for unrelated females as they were for older fecund females.

## Discussion

Our study examined the costs and benefits associated with two alternative reproductive tactics in banded mongooses, mate guarding and pestering. Contrary to our expectation that sneaking behaviour would represent a less costly reproductive tactic than mate guarding, we found that pestering males lost a similar percentage of body weight over the oestrus period as guarding males. Males who did not participate in pestering or mate guarding had no net weight loss. The weight loss associated with both pestering and mate guarding is likely driven by a combination of shared physical activity and the opportunity costs of reduced foraging [60,77]. We also found that pesterers had a clear siring disadvantage compared to mate guards and the probability of successful siring decreased with the proportion of rivals using the same tactic. These siring success results indicate that the fitness benefits of both pestering and mate guarding exhibited negative frequency dependent dynamics, consistent with alternative reproductive tactics in other species [13,24,30,33]. The adoption of pestering tactics by previously inactive males was consistently rarer than guarding, suggesting that pestering is not used as a stepping stone as males tend to remain reproductively inactive until they can gain mate-guarding roles. Pestering tactics were also only rarely adopted by guards, but pestering did increase for low weight and younger previous guards indicating pestering tactics are used as a back-up option when low RHP males are displaced.

The lack of evidence that pestering tactics are used as a ‘stepping stone’ to future guarding tactics is in contrast to their conditional use by growing males in other systems [30,43–45] before graduating to fighting tactics, e.g. Arctic char (*Salvelinus alpinus*) and pink salmon (*Oncorhynchus gorbuscha*)[78]. Instead, younger, low RHP male banded mongooses typically stay inactive until they can grow to gain a guarding position rather than adopt pestering tactics in the interim. One likely reason for the lack of pestering tactics being adopted by low RHP males is displacement by other pesterers. Displacement is suggested based on observations of pesterers chasing off other pesterers who approached the same guarded female, explaining the negative frequency dependent siring success found in this study. A small size may typically be beneficial for sneakers in other systems if it helps males remain undetected by fighters [44], but competition amongst pesterers for access to females would mean that any advantage of a small size may not be realised if males are displaced by higher RHP pesterers. Another reason for the lack of adoption of pestering tactics is the relatively high cost of pestering. Whereas using sneaky tactics can reduce reproductive costs compared to fighters in others systems [30], such as mortality costs in the two-spotted spider mite (*Tetranychus urticae Koch*) [48], pestering banded mongooses suffered similar weight loss costs compared to guarding males. These weight lost costs will be felt hardest by younger, low RHP males who may lack the condition required to sustain the energetic demands of reproductive activity compared to older, higher RHP males in the group [46,47]. Overall, low RHP male banded mongooses may accrue fewer benefits and incur higher costs of pestering than is the case for adopters of sneaky tactics in other systems, explaining their lack of adoption as an initial early opportunity for reproductive success on the way to guarding tactics in later life.

The weight loss costs associated with pestering could also lead to future fecundity costs. For example, in African striped mice (*Rhabdomys pumilio*) males that adopt roaming tactics suffer a delay in the weight gain needed to defend future harems, which can reduce life-time reproductive success [41,42]. Similarly, in banded mongooses, our transition matrices indicate that subordinates needed to reach a certain weight relative to other group members to transition into mate guarding tactics. Thus, weight loss by pesterers could hinder their growth and the acquisition of future profitable guarding roles, diminishing their lifetime reproductive fitness. In contrast, for males with a higher RHP pestering is unlikely to have an as determinantal impact on their future fecundity as they have more resources to maintain reproductive activity and remain competitive, making pestering a less costly back-up option. Given the unprofitability of pestering and mate guarding for young and low weight males, these individuals may gain from investing in helping instead [79,80].

While pestering tactics are rare, there is evidence to support their use as a back-up option if males are displaced from their position as a mate guard. The use of alternative reproductive tactics should be more common under higher competition. For example, under high population densities African striped mice (*Rhabdomys pumilio*) increasingly use philopatric inheritance strategies, as males using dispersing strategies have reduced success at gaining their own harems [56]. However, for banded mongooses there was a lack of evidence that groups with higher male sex skew, therefore higher competition, contribute to the adoption of pestering tactics by either previous inactive or guarding banded mongooses. Yet, since male-biased sex ratios are the norm for banded mongoose groups any potential effect of the number of potential rivals for each female may be overshadowed by RHP differences between individual competitors. Among previous guards, lower weight and younger males in the group had the highest probability to transition into pestering tactics, suggesting that pesterers were often previous low RHP guards that had been displaced. Additionally, although we did not find evidence that pestering is a ‘stepping stone’ tactic for previously inactive males to switch to as they grow, a small number of inactive males did transition into pestering tactics. This minority of males may be on the cusp of becoming mate guards since the transition to pestering and guarding tactics by inactive males both increased with RHP relative to other group members.

While pestering tactics involved substantial costs and were relatively rare within groups, we found that by adopting pestering tactics males removed restrictions on mate choice. Younger guards had lower RHP and were restricted to guarding younger females due to assortative mating dynamics while the oldest males guarded the oldest females [49]. Assortative guarding interactions align with previous results showing that the oldest male banded mongooses sire the majority of groups’ offspring through copulating with oldest most fecund females [61]. Pesterers instead interacted with females of age ranks older than would be expected based on assortative mating of mate guards, suggesting that freedom to choose mates in a pestering tactic improves access to fecund females. A chance to sire from more fecund older females may partially compensate for a reduced chance of siring success compared to more assured siring success guarding lower quality females, while also facilitating access to females in cases where all partners are guarded by higher RHP rivals.

Pestering tactics may serve as a mechanism for inbreeding avoidance, important for cooperative breeders due to the overall high relatedness between members of typically family groups [81]. We found both guards and pesterers displayed a preference for interacting with less closely related females in the group, aligning with previous findings indicating inbreeding avoidance among mate-guarding males in banded mongooses [65].This preference implies that low RHP guards do not face the same level of constraint in mate choice based on relatedness compared to mate choice based on a females age. However, our transition results suggest that lower RHP guards, who may only have had access closely related females, have the option to switch to pestering tactics instead of inbreeding or settling for reproductive inactivity. Restricted choice for unrelated females may be more common for lower RHP guards, which our transition results show is associated with guarding to pestering transitions. Flexible switching to pestering tactics may explain how guards and pesterers maintain a similar preference for less related females in the group, suggesting indirect evidence of the adoption of sneaky ARTs as a back-up option for inbreeding avoidance.

Females may benefit from the use of pestering tactics by males. Female mate choice can prevent less attractive males from exercising mate choice, compared to attractive males who maintain multiple partner options who can continue to exhibit choosiness [63,82,83]. In banded mongoose groups, males guard access to females but females can have some control over their partners by making attempts to escape their guard [58]. Although both sexes prefer to avoid close relatives as mates, male-biased sex ratios in groups mean females may often disagree with their guard over partner choice as females have a larger pool of partners to choose from. Additionally, as females are limited in the offspring they can produce, they should be choosier than males who can sire offspring from multiple litters [62,84]. Females may attempt to escape their guard to avoid inbreeding, evidenced by previous work showing that in cases of non-guard sires the successful sire is typically less related to the female than the failed mate guard [65]. Pesterers may act as a convenient alternative partner for females to avoid inbreeding in cases where females disagree with a guard’s advances. Even if the guards have copulated, any successful copulation with pesterers may allow potential cryptic female choice mechanisms to select the most compatible sperm [84,85], which may explain some of the skew towards less related sires found previously [65]. However, whether female choice mechanisms are present in banded mongooses is not currently known. Pestering tactics may benefit both males and females by expanding partner options that were previously restricted by guarding males and facilitating inbreeding avoidance.

Sneaky ARTs seem likely to be more widespread within social groups then is currently reported. It is common for social groups to include multiple mature males [86–88]. Subordinates can provide value to the group such as in offspring care [88–90]. In mammal societies, multiple male group-members can provide security to social groups by defending against outgroup usurping males [91] or in intergroup conflict [70]. These subordinate males may always have an incentive to steal copulations from higher RHP group members. Underreporting of sneaky ARTs may be because they are difficult to observe, oestrus periods are often fleeting and sneaking of copulations may happen out of sight. We benefited from an accumulation of observations over decades of study, which we have used to shed light on the strategy behind the adoption and use of sneaky ARTs and their potential role in inbreeding avoidance in banded mongooses. Where the challenges with the observation of sneaky tactics can be overcome, similar research into sneaky ARTs warrants expansion to other social systems.

## Conclusion

We did not find evidence that sneaky ARTs are used a stepping stone reproductive strategy in male banded mongooses. The costs and depreciating siring success of pestering may typically select for young, light subordinates to remain reproductively inactive until they have a RHP high enough to successfully adopt a mate guarding tactic. Instead, sneaky ARTs in male banded mongooses are rare and we suggest are used by a minority of males as back-up option if mate guarding positions are lost or fail to be secured. However, where pestering tactics are adopted, pesterers take advantage of reduced restrictions on mate choice by strategically interacting with the oldest, most fecund females in the group, suggesting that sneaky ARTs can be used by low RHP males to improve male mate choice options in social groups. We further suggest these sneaky ARTs may facilitate inbreeding avoidance for both males and females. Research into the strategic importance of sneaky ARTs within social groups is scarce, and warrants expansions to other social systems.

## Acknowledgements

Special thanks to Nicolas Fasel for help getting started using State transition models in JAGS. Additionally thanks to current and past members of our research group for helpful discussion and input. Additionally thanks to current and past members of our research group for helpful discussion and input.

