## Supplementary information for "Strategic use of male alternative reproductive tactics in cooperatively breeding banded mongoose groups"

**Supplementary material for alternative reproductive tactics in male banded mongooses**

**The relationship between age and weight in banded mongooses and weight inputation method**

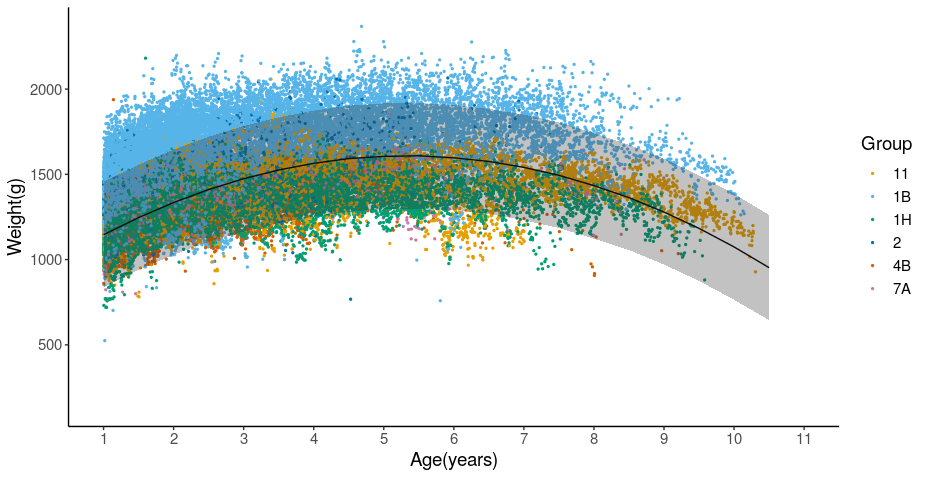

Supfig1: The relationship between age (years) and bodymass of males. Dots represent historically collected weights from 6 groups in the Mweya study population. Line and ribbon represent the output of a General Linear Model for age predicting weight with group as a random effect. Data show that weight increased with age until around 6 years of age, and then decreased.

**Supplementary methods**

*Weight inputation*

Weight is correlated with age in banded mongooses, and 93.75% of males (7/112) that required imputed weights had some history of weight collection allowing imputation to track individual weight trajectories using random slopes. Missing data was inputted using a linear mixed effect model fitted with quadratic age, random intercept (group id), and random slope on male id with quadratic age using the lme4 r package. The model was run using 48500 weights collected on the 314 males of known age above 180 days old collected since 2000. This included all males that required inputted weights. The quadratic term was justified to account for increases in age as individuals grew, and decreases in weight with senescence. This was validated by model comparisons with a linear model by likelihood ratio tests (Quadratic AIC = 28807, Linear AIC = 56608, Null AIC = 86050). Missing weights were extracted from the sampling distribution predicted from the above model simulated 10,000 times using the PredictInterval function (merTools r package).

**Supplementary results**

**Bayesian model output for all models in this study**

**Table S1**

| Summary of bayesian general linear models fitted in this study | | | | | | | | | |
| --- | --- | --- | --- | --- | --- | --- | --- | --- | --- |
| **Model** | **Term** | **effect** | **sd** | **2.5%** | **50.0%** | **97.5%** | **Rhat** | **f** | **Overlap0** |
| a | Constant(StaySub) | 0.03 | 0.19 | -0.32 | 0.03 | 0.4 | 1 | 0.56 | Yes |
|  | Constant(SubToPest) | -1.78 | 0.24 | -2.25 | -1.76 | -1.34 | 1 | 1 | - |
|  | Constant(SubToGuard) | 0.13 | 0.3 | -0.46 | | 0.13 | 0.73 | 1 | 0.68 |
|  | Constant(PestToSub) | -0.14 | 0.38 | -0.89 | -0.15 | 0.67 | 1.02 | 0.65 | Yes |
|  | Constant(StayPest) | -1.03 | 0.44 | -1.88 | -1.05 | -0.18 | 1 | 0.98 | - |
|  | Constant(PestToGuard) | 0.51 | 0.46 | -0.35 | 0.5 | 1.42 | 1.01 | 0.89 | Yes |
|  | Constant(GuardToSub) | -0.53 | 0.23 | -0.93 | -0.55 | -0.08 | 1 | 0.98 | - |
|  | Constant(GuardToPest) | -1.6 | 0.27 | -2.15 | -1.61 | -1.04 | 1 | 1 | - |
|  | Constant(StayGuard) | 0.8 | 0.27 | 0.27 | 0.8 | 1.3 | 1 | 1 | + |
|  | **AgeRank(StaySub)** | 0.15 | 0.02 | 0.11 | 0.15 | 0.19 | 1 | 1 | + |
|  | **AgeRank(SubToPest)** | 0.12 | 0.03 | 0.07 | 0.12 | 0.17 | 1 | 1 | + |
|  | **AgeRank(SubToGuard)** | -0.17 | 0.02 | -0.22 | -0.17 | -0.13 | 1 | 1 | - |
|  | **AgeRank(PestToSub)** | 0.12 | 0.05 | 0.03 | 0.12 | 0.21 | 1.03 | 0.99 | + |
|  | AgeRank(StayPest) | 0.09 | 0.06 | -0.02 | 0.09 | 0.2 | 1 | 0.94 | Yes |
|  | **AgeRank(PestToGuard)** | -0.16 | 0.06 | -0.28 | -0.16 | -0.05 | 1.01 | 1 | - |
|  | **AgeRank(GuardToSub)** | 0.2 | 0.03 | 0.14 | 0.2 | 0.27 | 1.01 | 1 | + |
|  | **AgeRank(GuardToPest)** | 0.15 | 0.04 | 0.08 | 0.15 | 0.23 | 1 | 1 | + |
|  | **AgeRank(StayGuard)** | -0.23 | 0.04 | -0.3 | -0.23 | -0.16 | 1 | 1 | - |
|  | **Weight(StaySub)** | -0.52 | 0.09 | -0.69 | -0.52 | -0.35 | 1 | 1 | - |
|  | Weight(SubToPest) | -0.15 | 0.13 | -0.39 | -0.15 | 0.1 | 1 | 0.88 | Yes |
|  | **Weight(SubToGuard)** | 0.57 | 0.09 | 0.38 | 0.57 | 0.74 | 1 | 1 | + |
|  | **Weight(PestToSub)** | -0.66 | 0.24 | -1.15 | -0.65 | -0.22 | 1 | 1 | - |
|  | **Weight(StayPest)** | -0.6 | 0.28 | -1.18 | -0.6 | -0.07 | 1 | 0.98 | - |
|  | **Weight(PestToGuard)** | 0.74 | 0.26 | 0.26 | 0.74 | 1.29 | 1.01 | 1 | + |
|  | **Weight(GuardToSub)** | -0.62 | 0.14 | -0.9 | -0.62 | -0.36 | 1 | 1 | - |
|  | **Weight(GuardToPest)** | -0.56 | 0.17 | -0.88 | -0.55 | -0.21 | 1 | 1 | - |
|  | **Weight(StayGuard)** | 0.72 | 0.15 | 0.44 | 0.72 | 1.02 | 1.01 | 1 | + |
|  | GroupSexRatio(StaySub) | -0.09 | 0.11 | -0.29 | -0.09 | 0.13 | 1.01 | 0.81 | Yes |
|  | GroupSexRatio(SubToPest) | -0.06 | 0.15 | -0.36 | -0.06 | 0.21 | 1 | 0.63 | Yes |
|  | GroupSexRatio(SubToGuard) | 0.13 | 0.13 | -0.11 | 0.13 | 0.38 | 1 | 0.83 | Yes |
|  | GroupSexRatio(PestToSub) | -0.04 | 0.23 | -0.47 | -0.04 | 0.42 | 1 | 0.57 | Yes |
|  | GroupSexRatio(StayPest) | -0.01 | 0.27 | -0.6 | 0 | 0.49 | 1 | 0.5 | Yes |
|  | GroupSexRatio(PestToGuard) | -0.07 | 0.32 | -0.66 | -0.07 | 0.56 | 1.01 | 0.59 | Yes |
|  | GroupSexRatio(GuardToSub) | 0.03 | 0.14 | -0.25 | 0.03 | 0.29 | 1 | 0.61 | Yes |
|  | GroupSexRatio(GuardToPest) | 0.22 | 0.16 | -0.13 | 0.22 | 0.53 | 1.01 | 0.92 | Yes |
|  | GroupSexRatio(StayGuard) | -0.07 | 0.17 | -0.4 | -0.06 | 0.25 | 1 | 0.66 | Yes |
|  | Random effect | Group | Male.id | Oestrus.event |  |  |  |  |  |
|  | sd | 0.0725 | 0.197 | 0.477 |  |  |  |  |  |
| b | Constant | -0.02 | 0.83 | -1.71 | -0.02 | 1.61 | 1 | 0.51 | Yes |
|  | **Guard(vs pesterer)** | 1.61 | 0.41 | 0.81 | 1.6 | 2.41 | 1 | 1 | + |
|  | **Competitors(n)** | -0.76 | 0.21 | -1.16 | -0.75 | -0.35 | 1 | 1 | - |
|  | **Competitors(shared tactic)** | -3.01 | 1.48 | -6.06 | -2.98 | -0.13 | 1 | 0.98 | - |
|  | Random effect | Female.id | Male.id | Oestrus.event |  |  |  |  |  |
|  | sd | 0.019 | 0.410 | 0.014 |  |  |  |  |  |
| c | Constant | -1.51 | 0.35 | -2.16 | -1.49 | -0.81 | 1 | 1 | - |
|  | **Subordinate** | 1.74 | 0.26 | 1.25 | 1.74 | 2.2 | 1 | 1 | + |
|  | **Pesterer** | -0.91 | 0.32 | -1.53 | -0.91 | -0.3 | 1 | 1 | - |
|  | **Guard** | -0.83 | 0.28 | -1.44 | -0.82 | -0.33 | 1 | 1 | - |
|  | Random effect | Group | Male.id | Oestrus.event |  |  |  |  |  |
|  | sd | 0.593 | 0.137 | 0.529 |  |  |  |  |  |
| d | Constant | -0.54 | 0.12 | -0.75 | -0.54 | -0.32 | 1.04 | 1 | - |
|  | **FemaleAgeRank** | -0.4 | 0.05 | -0.49 | -0.39 | -0.29 | 1.01 | 1 | - |
|  | **MaleAgeRank** | -0.07 | 0.04 | -0.15 | -0.07 | 0 | 1.02 | 0.97 | - |
|  | **Relatedness** | -0.63 | 0.18 | -0.96 | -0.63 | -0.26 | 1 | 1 | - |
|  | **Guard(vs pesterer)** | -0.12 | 0.06 | -0.23 | -0.12 | -0.01 | 1.04 | 0.99 | - |
|  | **Female:MaleAgeRank** | 0.21 | 0.04 | 0.13 | 0.21 | 0.29 | 1.01 | 1 | + |
|  | FemaleAgeRank:Guard* | 0.1 | 0.05 | -0.01 | 0.1 | 0.2 | 1.01 | 0.97 | +* |
|  | MaleAgeRank:Guard | 0.05 | 0.05 | -0.03 | 0.05 | 0.16 | 1.02 | 0.87 | Yes |
|  | **Female:MaleAgeRank:Guard** | 0.22 | 0.05 | 0.12 | 0.22 | 0.32 | 1 | 1 | + |
|  | **NumberOfFemales** | -0.13 | 0.01 | -0.16 | -0.13 | -0.11 | 1 | 1 | - |
|  | Random effect | Female.id | Male.id | Oestrus.event |  |  |  |  |  |
|  | sd | 0.449 | 0.0208 | 0.0121 |  |  |  |  |  |

Output table for all models in this study, fit using JAGS MCMC. Numbers (1-5) correspond to tactic transition (1: RHP, 2: Competition), siring probability (3), weight loss (4), and mate choice models (5). Means (effect), Credible intervals (0.025,0.975), and medians (50%) effects for each covariate are sampled from the untransformed posterior distribution of each model. f is the proportion of the posterior distribution with the same sign as the mean (f=0.5 for no effect, f=0.975+ is equivalent to p<0.05). Overlap 0 shows whether 0 overlaps with the range of 2.5% and 97.5% quantiles of the posterior distribution for each fitted parameter, and bold covariates are those that had a significant effect (no overlap). Asterisks (*) represent results where posteriors overlapped with 0 but by a marginal amount (f>0.95). Where there was no, or marginal, overlap, the direction of the effect is given in ovelap0. Rhat is a measure of chain convergence (<1.1) [88]. The standard deviation for each random effect fitted (Group, Female.id, Male.id, Oestrus.event) is given for each model.

**Weight change over an oestrus event given the reproductive tactic adopted by males**

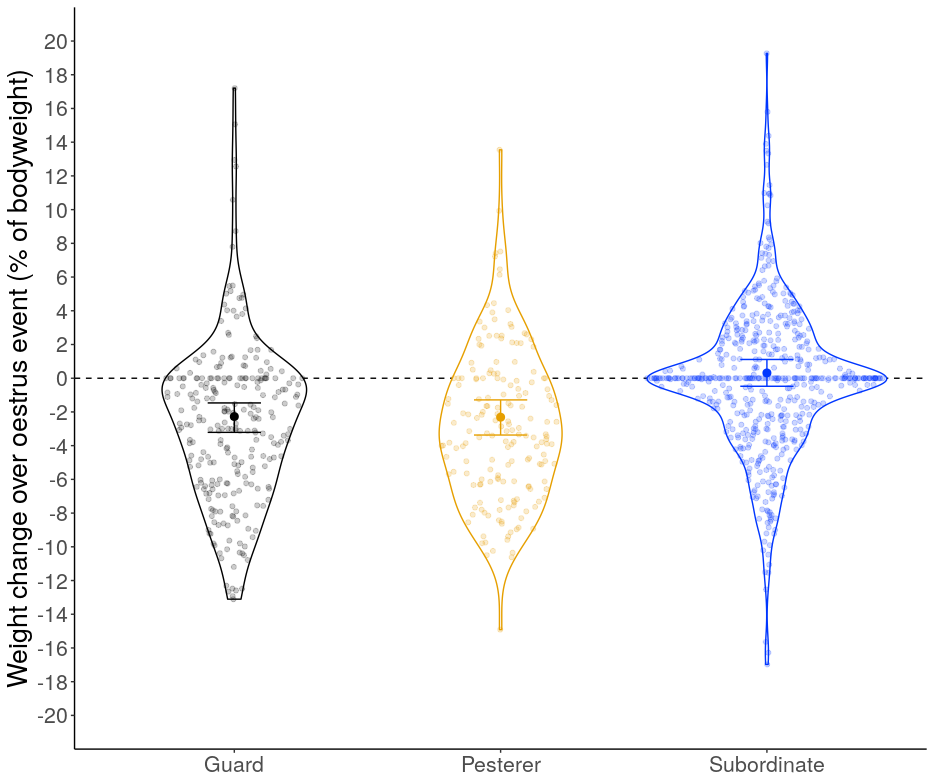

Figure S1: The effect of reproductive state of males on their weight loss over an oestrus event. Single bold points represent the modelled mean weight change, and error bars upper and lower credible intervals. Individual observed weight change is shown by shaded points, and their density distribution represented by a violin plot. The dashed lined represents neutral (0) weight change.

**Probability of male succesfully sires a females offspring**

Figure S2: The effect of competitor number and tactic adopted by a given male on their probability of siring offspring from the same female, given one of those competitors is a guard and the rest are pesterers. Mean posterior probabilities (points) and credible intervals (error bars) are shown. Raw success and failures to sire offspring by an individual male are displayed in text at the top, and as proportions as slim bars. For clarity, data are shown for cases where a female was guarded by a single male throughout the oestrus event. See SI for those rare cases where females were separately guarded by more than one unique male.

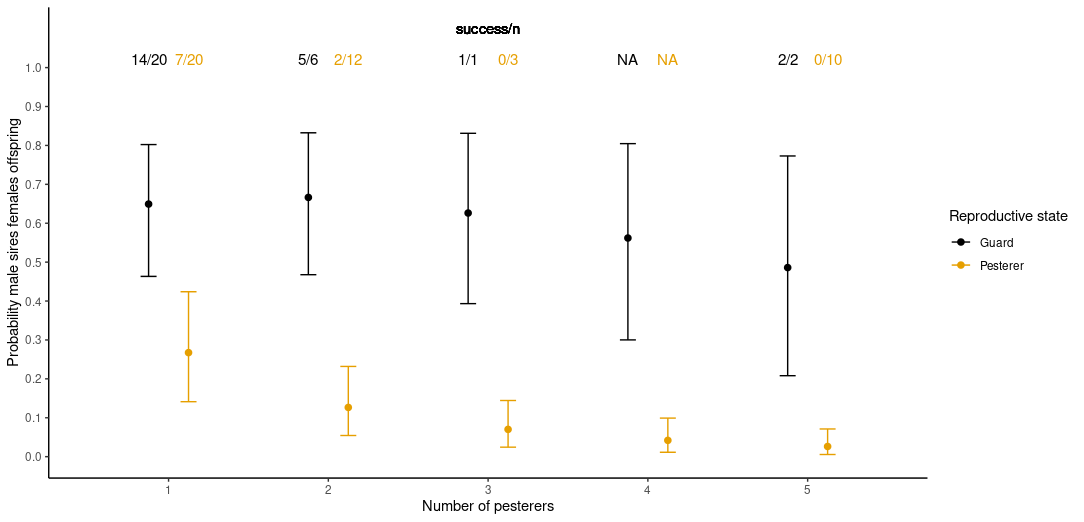

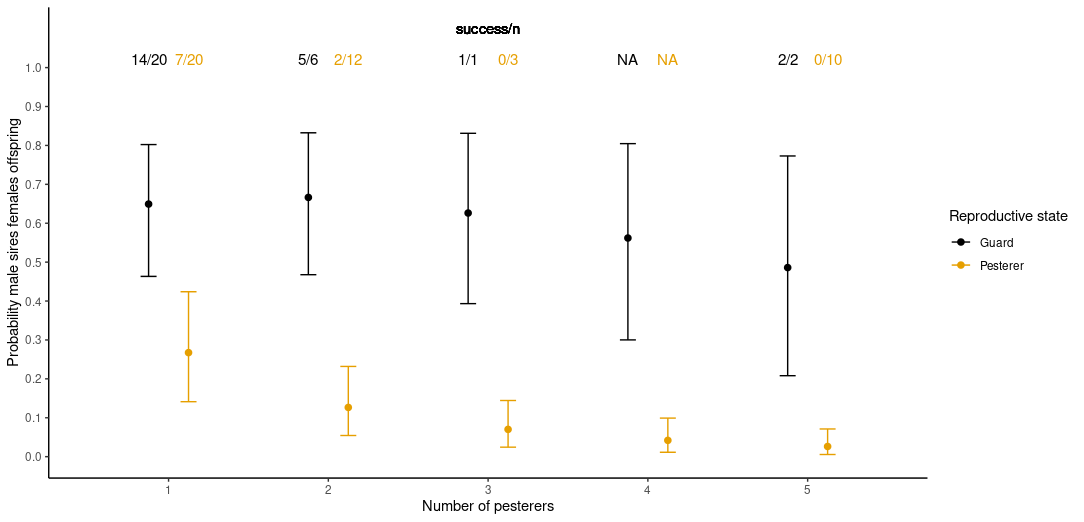

**The lack of an effect of competition on transitions between tactics**

There was no significant interaction between the effect of group adult sex ratio on age rank or group centered-weight on any of the transition probabilities. Group adult sex ratio had no significant effect on transitions from subordinate or pestering tactics (table S1), and simulated probabilities largely reflected the output of the null model (Figure 4a,b). There was no significant effect on the probability of keeping a guarding tactic or guard to subordinate transitions (table 1). Group adult sex ratio had no significant effect on guarding to pestering transitions (table 1: f=0.92), although a weak positive effect on guard to pestering transitions was found when age was fitted instead of age rank (supplementary, f=0.97), or when age rank and group centered weight were not fitted in the same model (supplementary, f=0.97).

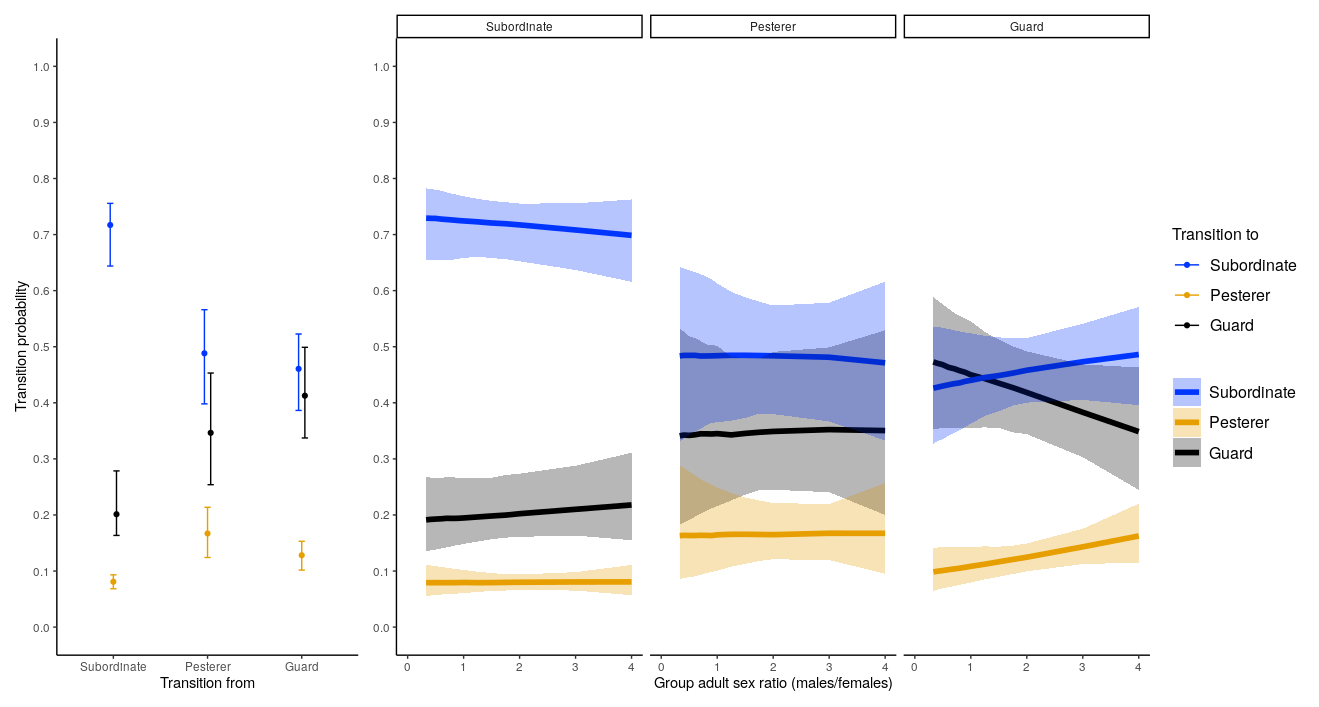

Figure S3: The effect of group adult sex ratio on between-oestrus event state transition probabilities. Lines represent the mean posterior probability and ribbons are 95% credible intervals. Each panel represents transition probabilities from the 3 distinct states. Black lines and ribbons correspond to transitions to guarding, orange transitions to pestering, and blue transitions to subordinate tactics. Panel (a) shows mean (points) and credible intervals (error bars) for all 9 transitions for reference based of a null model with no covariates.
